# Viewpoint-Dependence and Scene Context Effects Generalize to Depth Rotated 3D Objects

**DOI:** 10.1101/2022.11.15.516659

**Authors:** Aylin Kallmayer, Melissa L.-H. Võ, Dejan Draschkow

**Author notes:** Corresponding author contact information: Aylin Kallmayer, Department of Psychology, Scene Grammar Lab, Goethe University, Frankfurt, Germany, Mail.

## Abstract

Viewpoint effects on object recognition interact with object-scene consistency effects. While recognition of objects seen from “accidental” viewpoints (e.g., a cup from below) is typically impeded compared to processing of objects seen from canonical viewpoints (e.g., the string-side of a guitar), this effect is reduced by meaningful scene context information. In the present study we investigated if these findings established by using photographic images, generalise to 3D models of objects. Using 3D models further allowed us to probe a broad range of viewpoints and empirically establish accidental and canonical viewpoints. In Experiment 1, we presented 3D models of objects from six different viewpoints (0°, 60°, 120°, 180° 240°, 300°) in colour (1a) and grayscaled (1b) in a sequential matching task. Viewpoint had a significant effect on accuracy and response times. Based on the performance in Experiments 1a and 1b, we determined canonical (0°-rotation) and non-canonical (120°-rotation) viewpoints for the stimuli. In Experiment 2, participants again performed a sequential matching task, however now the objects were paired with scene backgrounds which could be either consistent (e.g., a cup in the kitchen) or inconsistent (e.g., a guitar in the bathroom) to the object. Viewpoint interacted significantly with scene consistency in that object recognition was less affected by viewpoint when consistent scene information was provided, compared to inconsistent information. Our results show that viewpoint-dependence and scene context effects generalize to depth rotated 3D objects. This supports the important role object-scene processing plays for object constancy.

## Introduction

Object recognition happens fast, automatic, and in most cases seems effortless to us. Since our environment is highly dynamic, especially when interacting with it, one and the same object will produce a range of different images on the retina. In fact, it is very unlikely that an object would produce the same retinal image twice due to changes in viewpoint, lighting, reflections, or viewing distance. Still, our visual system is able to flexibly transform this variable visual input in a way that object identity can successfully be read out from the resulting abstract representations in higher areas of visual cortex (see DiCarlo & Cox, 2007).

Whether object recognition is viewpoint-dependent (recognition performance is sensitive to changes in viewpoints as indicated by accuracy and response-time (RT) data) or viewpoint-invariant (recognition performance is largely unaffected by changes in viewpoint) has been a debated topic (Biederman & Gerhardstein, 1993; Bülthoff & Edelman, 1992; Burgund & Marsolek, 2000; Charles Leek & Johnston, 2006; Edelman, 1995; Graf, 2006; Hayward, 2003; Hayward & Tarr, 1997; Jolicoeur, 1990; Leek et al., 2007; Lowe, 1987; Marr et al., 1978; Ratan Murty & Arun, 2015; Stankiewicz, 2002; Tarr & Bülthoff, 1995; Tarr & Pinker, 1989; Wilson & Farah, 2003). Since the early debates, there has been overwhelming consensus that object recognition is neither solely viewpoint-dependent nor solely viewpoint-invariant and that evidence for both can be observed depending on experimental task and stimuli (Foster & Gilson, 2002; Hamm & McMullen, 1998; Jolicoeur, 1990; Leek et al., 2007; Ratan Murty & Arun, 2015; Sastyin et al., 2015; Stankiewicz, 2002; Vanrie et al., 2002).

Past research has made great advances towards understanding the mechanisms that underly invariant object recognition, when objects are presented in isolation (i.e., DiCarlo & Cox, 2007). More recently, however, researchers have started to investigate the viewpoint problem in the context of object-scene processing. Object recognition rarely occurs in isolation where the only available information are the objects’ features. In our everyday lives, we encounter objects within certain contexts, which provides us with a pool of complex visual and multimodal information that is integrated during object recognition. Past research has shown that context facilitates object recognition (Biederman et al., 1982; Oliva & Torralba, 2007; for a recent review see Lauer et al., 2021). Evidence from behavioral as well as neurophysiological studies (e.g., Brandman & Peelen, 2017) suggest an interactive processing of objects and scenes. For instance, objects placed in semantically consistent contexts are recognized faster and more accurately, often referred to as the *scene-consistency effect* (Davenport & Potter, 2004; Palmer, 1975). Accordingly, models of object recognition have been updated to incorporate the integration of contextual information (Bar, 2004). Further, frameworks incorporating object-scene and object-object relations (e.g., the so-called *scene-grammar)* describe a set of internalized rules based on regularities found in real-world scenes that facilitate scene and object perception and guide our attention during different visual cognitive tasks (Draschkow & Võ, 2017; Josephs et al., 2016; Võ, 2021; Võ et al., 2019; Võ & Henderson, 2009; Võ & Wolfe, 2013a, 2013b).

Sastyin and clleagues (2015) conducted a series of experiments investigating the interaction between viewpoint and scene-consistency on object and scene recognition. They used photographic images of objects shown from canonical and accidental viewpoints and paired them with consistent or inconsistent scenes. They found a significant interaction between viewpoint and consistency where the viewpoint effect was weaker when consistent scene information was provided. From this they concluded that object recognition relied more on context information if the object was presented from an accidental viewpoint.

Here, in order to increase the external validity of these findings (Draschkow, 2022), we aimed to generalize the insights from 2D photographic images to 3D models of objects (Biederman & Gerhardstein, 1993; Gauthier et al., 2002; Logothetis et al., 1994; Poggio & Edelman, 1990; Zisserman et al., 1995). Recent work using 3D immersive environments has highlighted the importance of studying vision under more naturalistic constraints in order to investigate cognitive processes in the context of natural behavior (Draschkow et al., 2021; Helbing et al., 2020, 2022; Kristjánsson & Draschkow, 2021). An additional benefit of using 3D models is that we could probe a broad range of viewpoints and empirically establish accidental and canonical viewpoints, allowing for a broader representation of the viewpoints we encounter in our natural environment.

In the present study, we conducted three behavioral experiments. In our first two experiments, (Experiment 1a and 1b) we presented 3D models of real-world objects from six different angles (0°, 60°, 180°, 120°, 240°, 300°) rotated around the pitch axis in a word-picture verification task. Because rotating the objects around the pitch axis results in highly atypical viewpoints, we expected to find viewpoint-dependent recognition indicated by lower accuracy and slower RTs. In Experiment 1b, we wanted to replicate Experiment 1a with grayscale versions of the images, expecting similar effects of viewpoint as for Experiment 1a (Hayward & Williams, 2000). Experiments 1a and 1b also served to identify viewpoints which produced highest (canonical) and lowest (non-canonical) recognition performance which we then used in Experiment 2.

In Experiment 2, we paired 3D objects presented in canonical (0° rotation) and non-canonical (120° rotation) viewpoints with semantically consistent and inconsistent scenes. Our aim was to test if viewpoint-dependence and object-scene processing effects (Sastyin et al., 2015) generalize to depth rotated 3D models of objects.

## General Method

### Participants

Participants were recruited at Goethe-University Frankfurt am Main. The sample consisted of 12 participants who completed Experiment 1a (6 women, *M* = 23.92, range = 19–29), 12 different participants who completed Experiment 1b (8 women, *M* = 19, range = 18–22), and another set of 32 participants who completed Experiment 2 (25 women, *M* = 24.28, range = 18–51). The sample size of Experiment 2 was a priori chosen to be higher compared to previous studies which found reliable effects across multiple experiments with 20 participants (e.g., Sastyin et al., 2015). In Experiment 1a, all except for six participants were psychology students that were compensated with course credits, while the remaining participants volunteered for the experiment without any compensation. All had normal or corrected-to-normal vision, were native German speakers, and were unfamiliar with the stimulus materials. Written informed consent was obtained before participation, data collection and analysis were carried out according to guidelines approved by the Human Research Ethics Committee of the Goethe University Frankfurt.

### Stimulus Material

For Experiment 1a and Experiment 1b, we collected 100 3D models of objects from a broad range of categories such as furniture, foods, vehicles, plants, and electrical devices. Eighty-two of the 3D models were purchased from CG Axis Complete packages I, II, III, and V, 18 additional models were obtained free of charge from sources like TurboSquid and free3D. Each model was rotated around its pitch axis by 0°, 60°, 120°, 180°, 240°, and 300° degrees and sized to fit a 60cm x 60cm x 60cm box using the free 3D animation software Blender. A snapshot from each angle was systematically recorded in front of a gray background using the virtual reality software Vizward5 to create our final stimulus set of 600 images. Additionally, we created grey-scaled versions of these images for Experiment 1b using the GrayscaleEffect function in Vizard5 (https://docs.worldviz.com/vizard/latest/postprocess_color.htm).

For Experiment 2, we used the same 3D models as in Experiment 1 adding an additional 56 models collected from the CGAxis packages, resulting in a total of 156 models. Instead of creating snapshots of all six angles, we chose the two viewpoints that had previously produced the highest (canonical viewpoint, 0°) and lowest (non-canonical viewpoint, 120°) recognition performance averaged over Experiment 1a and Experiment 1b. We gray-scaled the images using the method described above.

Additionally, we collected 312 photographic images of scenes, one consistent and one inconsistent scene for each object. We defined a consistent scene as one in which we would expect the object to appear naturally. In both cases, the target object was not present in the scene. Most of the photographs were obtained from the SCEGRAM database (Öhlschläger & Võ, 2017) as well as from Google images.

### Procedure

To investigate the speed and accuracy of object recognition, while keeping the procedure comparable with previous studies, a word-picture verification task was employed for all experiments (Figure 1). Participants were instructed on screen as well as through standardized verbal instructions to decide as quickly and accurately as possible whether the object on screen matched the basic level category label presented to them at the beginning of the trial using a corresponding “match” or “mismatch” key. Participants were not made aware of the different viewpoint conditions beforehand. Each experiment consisted of three practice trials during which the instructor stayed in the room with the participant. More detailed procedure and trial sequences will be described in the individual Procedure sections of each experiment. Experiments 1a and 1b lasted approximately 30 minutes, Experiment 2 lasted approx. 12 minutes.

**Figure 1.**
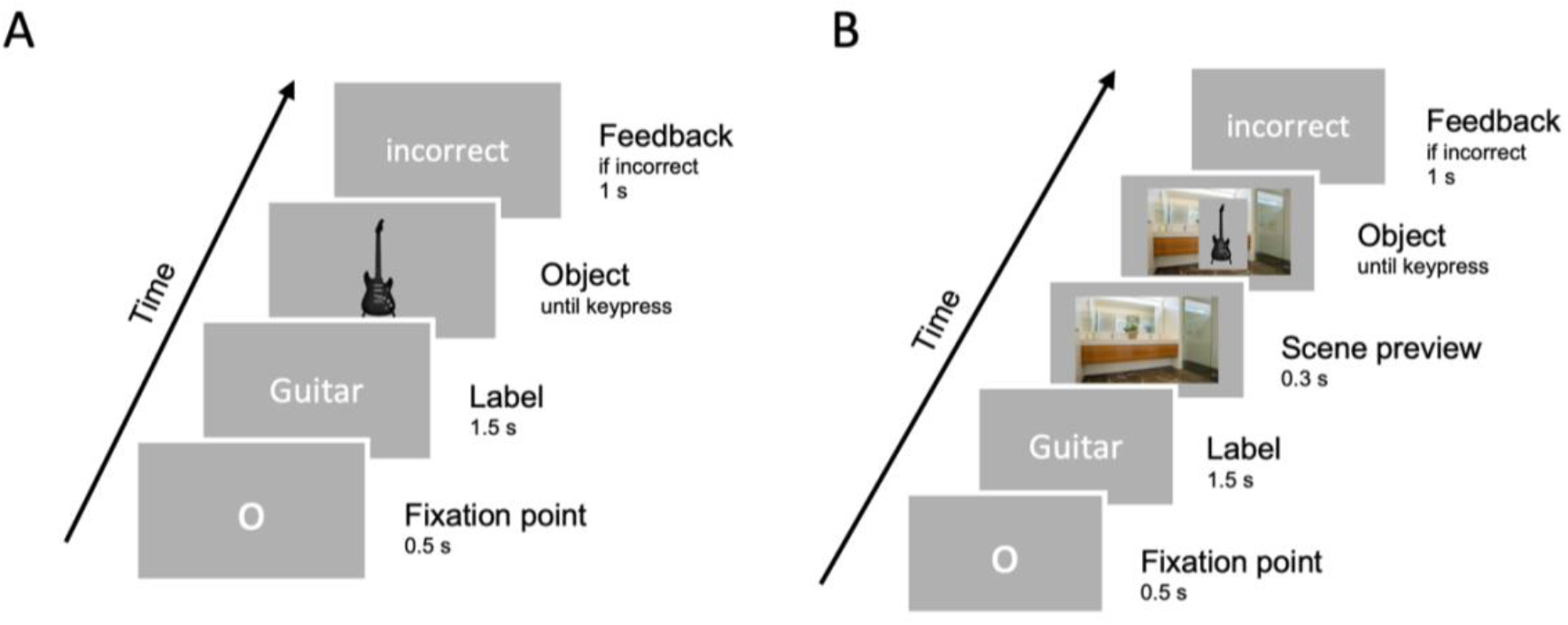
Trial procedures for the matching task in Experiment 1a and 1b (A) and Experiment 2 (B). The object was presented in colour in Experiment 1a and greyscaled in Experiment 1b. Note that the depicted labels are in English for visualization purpose. Feedback was only provided in case of incorrect responses.

### Design

Experiments 1a and 1b consisted of six blocks with 100 trials each. In each block, the object was presented from a different angle (0°, 60°, 120°, 180°, 240°, 300°) chosen randomly and counterbalanced between participants. The order of objects within each block was randomized. Each object appeared three times in the match condition (object image matched basic level category label) and three times in the mismatch condition (object image did not match basic level category label), randomized between blocks.

In the mismatch condition, the basic level category label stemmed from a different superordinate category than the object image (e.g., the label “chair” as part of the superordinate category “furniture” was paired with an image of a “car” as part of the superordinate category “vehicle”).

Because there was no effect of viewpoint in the mismatch condition in Experiment 1a and 1b, most trials in Experiment 2 were match trials (N = 120) with 23% mismatch trials (N = 36) that were later excluded from analysis. In Experiment 2, each object was presented to each participant once, and we counterbalanced consistency (consistent vs. inconsistent) and viewpoint (canonical vs. non-canonical) between participants.

### Data Analysis

In Experiments 1a and 1b, we were interested in the effects of viewpoint (how far the object was rotated away from its canonical 0° angle) and match (whether the object matched the basic level category label as part of the experimental design) on reaction times (time between the onset of the object image and keypress response) and accuracy. In Experiment 2, we were interested in the interaction between viewpoint (canonical versus non-canonical viewpoint), and scene consistency (consistent scene versus inconsistent scene) on reaction times and accuracy.

Raw data was pre-processed and analysed using R (R Core Team, 2021). Objects that produced accuracy ratings that deviated more than 2.5 SD from the mean (computed for each condition separately) were excluded from analysis. Based on this, we excluded four objects in Experiment 1a, one in Experiment 1b, and two in Experiment 2. We based our reaction time analysis on correctly matched trials only (percent trials removed: Experiment 1a = 4.45%, Experiment 1b = 10.16%, Experiment 2 = 8.55%).

In our data analysis, we employed (generalized) linear mixed-effects models ((G)LMMs) using the lme4 package (Bates et al., 2015). We chose this approach because of its potential advantages over analysis of variance (ANOVA) as it allows us to simultaneously estimate by-participant and by-stimulus variance (Baayen et al., 2008; Bates et al., 2014; Kliegl et al., 2011). The random effects structure of each model was determined using a drop-one procedure starting with the full model including by-participant and by-stimulus varying intercepts and slopes for the main effects in our design. We then subsequently removed random slopes that did not contribute significantly to the goodness of fit as determined by likelihood ratio tests. This allowed us to avoid overparameterization and produce converging models that are supported by the data. Details about the individual analysis and models are described in the Data Analyses sections of each experiment. For each GLMM we report β regression coefficients together with the *z* statistic and apply a two-tailed 5% error criterion for significance testing. *P*-values for the binary accuracy variable are based on asymptotic Wald tests. Additionally, reaction times were transformed following the Box-Cox procedure (Box & Cox, 1964) to correct for deviation from normality as to better meet LMM assumptions (see individual Data Analysis sections for further details). For the LMMs regression coefficients are reported with the t-statistic and p-values were calculated with the lmerTest package (Kuznetsova et al., 2017). We defined sum contrasts for match (match vs. mismatch), and consistency (consistent vs. inconsistent) where slope coefficients represent differences between factor levels and the intercept is equal to the grand mean.

We used the ggplot2 package (Wickham, 2016) for graphics and emmeans (Lenth, 2022) for post-hoc comparisons. Data and code are openly available at https://github.com/aylinsgl/2022-Viewpoint_and_Context.

### Apparatus

All experimental sessions were carried out in the same six experimental cabins of the department of psychology at Goethe-University Frankfurt am Main, containing the same experimental set up (computers running OS Windows 10). Stimulus presentation, response-times (RT) and accuracy were systematically controlled and recorded by OpenSesame (Mathôt et al., 2012), presented on a 19-in monitor (resolution = 1680 × 1050, refresh rate = 60 Hz, viewing distance = approx. 65 cm, subtending approx. 11.13 °× 9.28° of visual angle for the object images and approx. 19° × 15.84° of visual angle for the background images).

## Experiment 1a & 1b

In Experiments 1a and 1b, we investigated the effect of viewpoint on object recognition RT and accuracy using 3D models of objects rotated around the pitch axis (0°, 60°, 120°, 180°, 240°, 300°). The only difference between the experiments was that 3D models were presented either in color (Experiment 1a) or a grayscale version of the model was used (Experiment 1b). Participants had to indicate whether the object matched the previously presented basic level category label.

### Procedure

Participants were presented with a fixation point in the middle of the screen followed by a basic level object category label (in German, font: Droid Sans Mono; font size: 26; color: black). This was followed by the target object presented in the middle of the screen, which could either match or mismatch the label, until the participant gave a response (Figure 1A). Participants were given feedback on screen if their answer was incorrect. The next trial automatically started with a new fixation point.

### Data Analysis

After data preprocessing, we employed a binomial GLMM to examine the effects of viewpoint and match on accuracy. As fixed effects we included viewpoint (0°, 60°, 120°, 180°, 240°, 300°) as a first and second-degree polynomial, the match vs mismatch comparison, and the interactions between these terms. The second-degree polynomial viewpoint term was added as we expected viewpoint to affect recognition in a non-linear manner (symmetry around 180°). Our final model included random intercepts for participants and stimuli, as well as a by-stimuli random slope for the match vs. mismatch effect for Experiment 1a, and random intercepts for participants and stimuli, as well as a by-stimuli and by-participant random slope for the match effect for Experiment 1b.

Based on the power coefficient output of the Box-Cox procedure (λ = 0.22), RTs were log-transformed. We employed the same fixed effects structure for the RT-LMMs as for the accuracy-GLMMs. As random effects, we entered random intercepts for participants and stimuli, as well as by-participant and by-stimuli random slopes for the effect of match for Experiment 1a and 1b.

### Results

#### Accuracy

The average accuracy in Experiment 1a was quite high (*M* = 0.95, *SD* = 0.21) and slightly lower in Experiment 1b (*M* = 0.9, *SD* = 0.3). In line with our hypothesis, the GLMM yielded a significant main effect for the second-degree polynomial viewpoint term in both experiments (Experiment 1a: β = 16.67, *SD* = 5.61, *z* = 2.97, *p* = 0.003; Experiment 1b: β = 18.82, *SE* = 3.79, *z* = 4.97, *p* < 0.001), meaning that the effect of viewpoint on accuracy can be well described by a quadratic function (Figure 2A and 2C). There was also a significant interaction between the second-degree polynomial of viewpoint and the match condition in both experiments, Experiment 1a: β = 23.62, *SE* = 5.69, *z* = 4.15,*p* < 0.001; Experiment 1b: β = 15.23, *SE* = 3.82, *z* = 3.98, *p* < 0.001. Comparing the viewpoint trend for the match and mismatch conditions, we found that the second-degree viewpoint trend was significant in the match condition (Experiment 1a: β = 0.19, *SE* = 0.03, *CI*95% = [0.13, 0.25]; Experiment 1b: β = 0.16, *SE* = 0.02, *CI*95% = [0.12, 0.21), but not in the mismatch condition, Experiment 1a: β = −0.03, *SE* = 0.04, *CI*95% = [-0.12, 0.05]; Experiment 1b: β = −0.02, *SE* = 0.03, *CI*95% = [-0.04, 0.07].

**Figure2.**
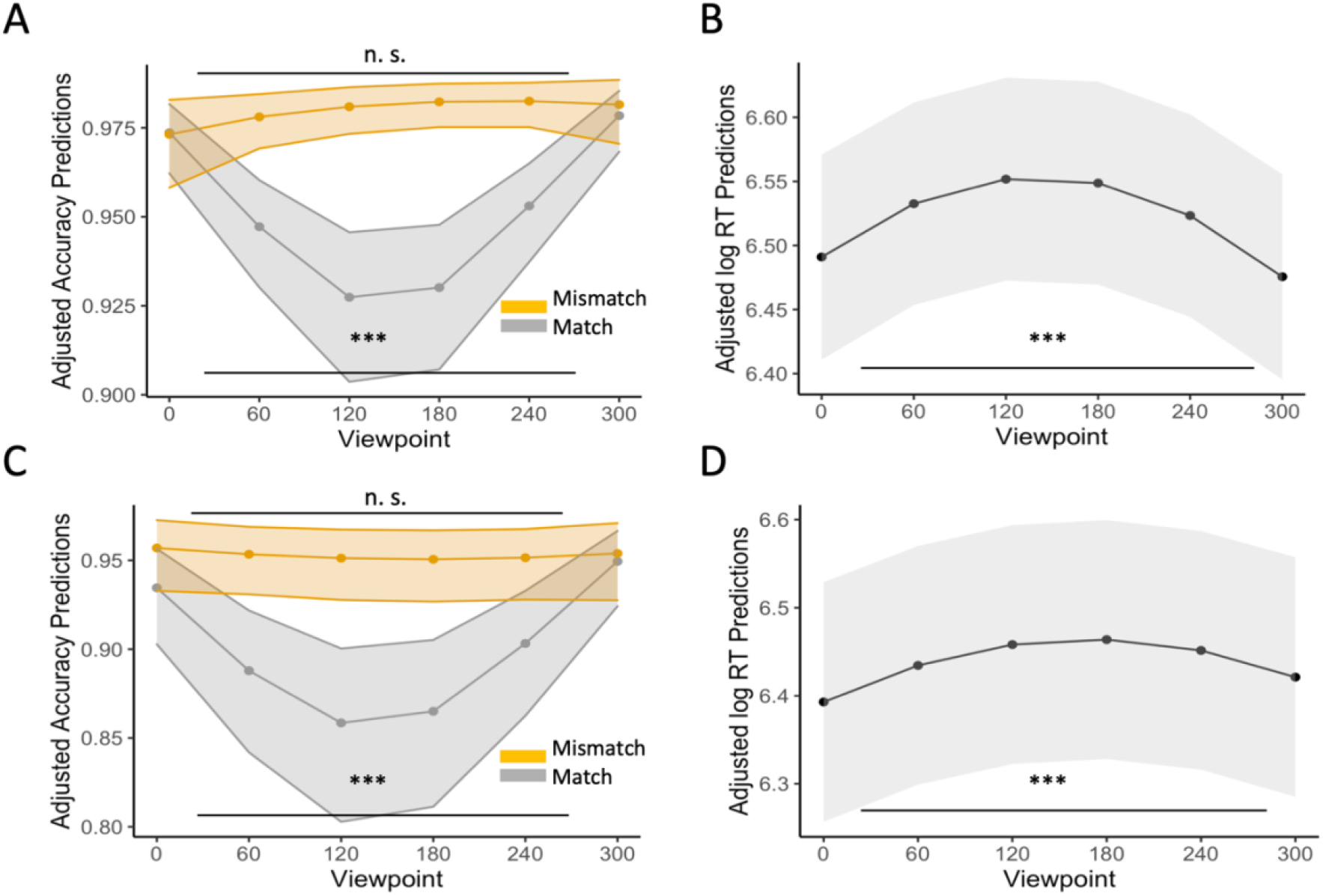
Partial effect plots of the interactions of viewpoint (0°, 60°, 120°, 180° 240°, 300°) and match (match vs. mismatch) on accuracy for Experiment 1a (coloured; A), and Experiment 1b (greyscaled; C), and the effect of viewpoint on RT for Experiment.

#### Response-times (RT)

Participants were slightly faster on average in Experiment 1b (*M* = 685 ms, *SD* = 358 ms) than Experiment 1a (*M* = 738 ms, *SD* = 299 ms). In line with our hypothesis, the LMM revealed a significant main effect for the second-degree polynomial viewpoint term in both experiments, Experiment 1a: β = −2.2, *SE* = 0.29, *t* = −7.48, *p* < 0.001; Experiment 1b: β = −1.42, *SE* = 0.29, *t* = −4.99, *p* < 0.001 (Figure 2B and 2D). In both experiments there was no interaction between viewpoint and match, Experiment 1a: β = −0.12, *SE* = 0.29, *t* = −0.4, *p* = 0.69; Experiment 1b: β = −0.38, *SE* = 0.29, *t* = −1.34, *p* = 0.18.

## Discussion

In Experiment 1a, we found viewpoint-dependent object recognition for objects rotated around the pitch axis. This effect can best be described by a quadratic curve that approximates symmetry around 120° rotation. We also found that in our sequential matching task, only the match condition produced viewpoint-dependent behavior, while mismatch trials seemed unaffected by viewpoint. Finding a mismatch might rely more on the analysis of global, viewpoint-invariant features, whereas matching might be more dependent on the analysis of local, viewpoint-dependent features (e.g., Jolicoeur, 1990a) (e.g., deciding a shape is not a car might require less viewpoint-dependent information than identifying the shape as a chair). In Experiment 1b, we were able to replicate our results from Experiment 1a. Grayscaling the images seemed to have made the overall task slightly more difficult while still producing similarly viewpoint-dependent behavior. The canonical (0°) and non-canonical (120°) viewpoints we used in Experiment 2 represented viewpoints that produced the best and worst recognition performance derived from average accuracy ratings obtained from Experiment 1a and 1b.

## Experiment 2

In Experiment 2, we paired canonical (0°) and non-canonical (120°) viewpoints with consistent and inconsistent scene contexts. We were specifically interested in the interaction between viewpoint and consistency with the expectation that meaningful scene context information would reduce the effect of viewpoint on object recognition.

### Procedure

In Experiment 2, we used the same word-picture verification task as in Experiments 1a and 1b (Figure 1B). Scene context was provided by first previewing the consistent or inconsistent scene for 300ms and then overlaying the target object on top of the scene background until a response was given.

### Data Analysis

For both the accuracy-GLMM and response time (RT) LMM we entered interaction terms between viewpoint and consistency as fixed effects. The GLMM included random intercepts for participants and stimuli, as well as a by-stimuli random slope for the effect of viewpoint. Response time data was log transformed.

For the RT-LMM we had random intercepts for participants and stimuli, and a by-participant random slope for the effect of viewpoint and by-stimuli random slopes for the effects of viewpoint and consistency.

### Results

#### Accuracy

Accuracy was significantly higher for canonical viewpoints than for non-canonical viewpoints as revealed by the GLMM (β = 0.68, *SE* = 0.14, *z* = 4.82, *p* < 0.001) but there was no significant main effect for consistency, β = 0.06, *SE* = 0.07, *z* = 0.75, *p* = 0.45. Critically, there was a significant interaction between viewpoint and consistency, β = −0.21, *SE* = 0.07, *z* = −2.84, *p* = 0.004 (Figure 3A). Post-hoc interaction contrasts revealed that the viewpoint-dependence effect was significantly stronger in the inconsistent scene condition compared to the consistent scene condition, β = −0.84, *SE* = 0.3, *z* = −2.84, *p* = 0.005. This is in line with our hypothesis that providing meaningful scene context can reduce the effects of viewpoint on object recognition. Additionally, the scene-consistency effect was only significant in the non-canonical condition (β = 0.53, *SE* = 0.15, *z* = 3.45,*p* < 0.001), but not in the canonical condition, β = −0.31, *SE* = 0.25, *z* = −1.22, *p* = 0.22.

**Figure 3.**
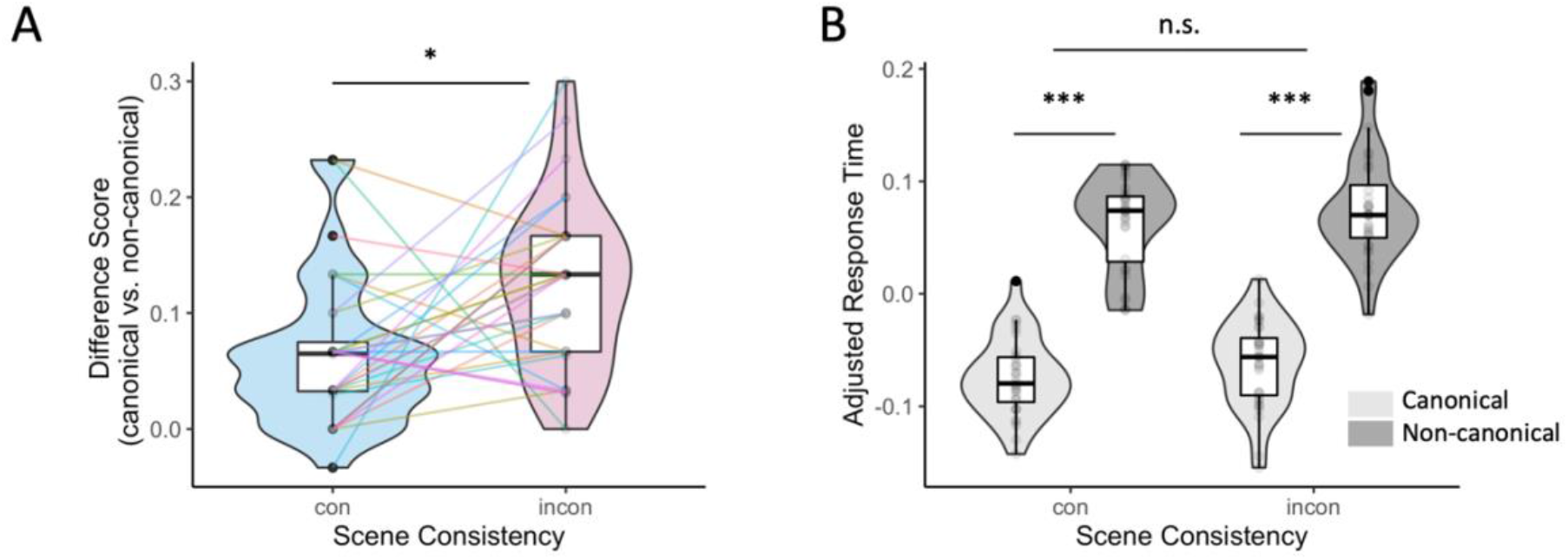
Experiment 2 accuracy difference scores per participant (canonical vs. non-canonical) for consistent and inconsistent scene backgrounds (A). Adjusted response times (B) were obtained with the remef package (Hohenstein & Kliegl, 2021). *p < .05. ***p < .001.

#### Response-Times (RT)

The LMM yielded a significant main effect for viewpoint (β = −0.07, *SE* = 0.01, *t* = −7.26, *p* < 0.001), where RTs were faster for canonical (*M* = 558ms, *SD* = 255ms) than for non-canonical viewpoints (*M* = 645 ms, *SD* = 333 ms) (Figure 3B). There was no significant interaction between viewpoint and consistency, β = 0.004, *SE* = 0.005, *t* = 0.83, *p* = 0.41.

### Discussion

In general, object recognition accuracy was viewpoint dependent, however, there was a significant interaction between viewpoint and consistency. In line with our hypothesis, the viewpoint effect was significantly weaker for consistent scenes and the scene consistency effect was only observed for non-canonical viewpoints (Figure 3A). Non-canonical viewpoints were recognized significantly slower than canonical viewpoints. However, this was unaffected by scene consistency.

## General Discussion

In the present study, we investigated how scene context information modulates viewpoint-dependent object recognition using 3D models of everyday objects. While providing meaningful context did not eradicate the viewpoint effect fully, it significantly reduced recognition accuracy costs. In line with previous findings (Sastyin et al., 2015) this supports a model of object recognition that incorporates context (e.g., Bar, 2004) while dynamically adapting to the amount of available information based not only on visual features of the object (Burgund & Marsolek, 2000; Hayward & Tarr, 1997; Jolicoeur, 1990), but also context. It further motivates models of object constancy - the visual system’s ability to produce representations that are robust to changes in e.g., viewpoint or lighting (e.g., DiCarlo & Cox, 2007) – that efficiently integrate contextual information and can lead to both viewpoint-dependent and invariant behavior based on available information and the task at hand.

A key component of the present study was to generalize previous findings on object-scene processing effects and viewpoint-dependence to depth rotated 3D objects. We want to highlight the importance of generalizing findings from traditional 2D settings to more naturalistic settings and stimuli. Kristjánsson and Draschkow., (2021) have shown very illustratively for a variety of phenomena that given more naturalistic constraints, a system is able to circumvent e.g., capacity limits by drawing on the rich visual experience of natural environments. While we did not use fully immersive environments, using 3D models offers a more realistic encounter of everyday objects and therefore a more precise measure of viewpoint-dependence in real-world object recognition. It should be noted, however, that there is a trade-off between naturalistic *looking* stimuli (i.e., photographs) and stimuli that more precisely capture naturalistic properties (i.e., 3D structure of objects from different viewpoints) in a highly controlled manner while not *looking* as naturalistic. Here, we opted for providing more naturalistic 3D properties of the displayed objects.

From the present study it is unclear what kind of information contained in the scenes was responsible for reducing the viewpoint costs. Rapidly accessed global information such as the gist of the scene (Oliva & Torralba, 2007) could be the main factor. At the same time, more local information such as the detection and recognition of certain objects in the scene preview could provide information about related possible target objects based on internalized scene-object and object-object regularities (Võ et al., 2019). Revealing the time course of when what kind of contextual information is integrated to buffer viewpoint effects would provide new insights into how the visual system so effortlessly achieves invariant object recognition.

Varying what information is presented during the task (i.e., providing meaningful context vs. showing objects in isolation) is one way to probe the visual system’s ability to overcome processing limitations in viewpoint-dependent object recognition. Alternatively, one could keep the visual input constant but vary the level at which participants have to perform the matching task (Hamm & McMullen, 1998). If there are object representations that contain more or less viewpoint-dependent or invariant information how does this interact with the integration of contextual information in the form of scene context?

Finally, we would like to address that on average performance was high in the matching task throughout all our experiments. These ceiling effects are probably due to the type of task we chose - different from the tasks usually employed to study scene consistency effects (Davenport & Potter, 2004; Sastyin et al., 2015). Despite these differences in difficulty, we were able to demonstrate a significant reduction in viewpoint costs by providing meaningful scene context.

Past research has made strong advances towards understanding the computations that underly invariant object recognition (DiCarlo & Cox, 2007). Understanding these mechanisms in isolation is key to understanding object recognition in general. We argue that understanding how the visual system is able to make use of richly structured naturalistic environments to circumvent computational bottlenecks will ultimately lead to better, more robust models of object recognition and inspire approaches in fields such as computer vision (e.g., Bomatter et al., 2021).

To conclude, in the present study we built upon previous findings on object-scene processing and viewpoint dependence by generalizing these effects to depth rotated 3D objects. We highlight the importance of testing capacity limits of object recognition in more naturalistic frameworks in order to build more robust and flexible models and move towards a better understanding of vision under naturalistic constraints.

## Acknowledgements

This work was funded by the Deutsche Forschungsgemeinschaft (DFG, German Research Foundation)—project number 222641018—SFB/TRR 135, sub-project C7 to M.L.-H.V., and by the Main-Campus-doctus scholarship of the Stiftung Polytechnische Gesellschaft Frankfurt a. M. to A.K.

The Wellcome Centre for Integrative Neuroimaging is supported by core funding from the Wellcome Trust (203139/Z/16/Z). The work is supported by the NIHR Oxford Health Biomedical Research Centre. The funders had no role in the decision to publish or in the preparation of the manuscript.

